# Temporal constraints on the neural signatures of narrative processing

**DOI:** 10.64898/2026.07.09.737443

**Authors:** Aline-Priscillia Messi, Abir Bhuyain, Liina Pylkkanen

**Author notes:** Corresponding author: Aline-Priscillia Messi 10 Washington Place New York University New York, NY 10003.

## Abstract

How the brain constructs meaning across extended contexts remains poorly understood. While neural responses to words and sentences are well characterized, much less is known about the brain mechanisms supporting narrative comprehension. Sentence-level studies suggest that neural activation increases as word meanings are integrated into sentence meaning. At the discourse level, theories propose that narratives depend on situation models, possibly engaging networks beyond core language regions, including the default mode network. Because narrative comprehension unfolds over longer timescales, processing time may be a bottleneck. In this MEG study, we tested how representation size and presentation rate shape neural responses by varying linguistic structure (words, sentences, stories) and the speed of visual text in 1–4-word chunks. We found an early bilateral story effect in visual cortex, followed by a spatiotemporal progression of activity along the temporal lobes that culminated in a three-way contrast among word lists, sentence lists, and stories. Faster presentation altered this pattern: the left-lateralized story effect disappeared, and the right-lateralized effect became more spatially restricted. Under Fast presentation, significant effects were limited to left lateral language cortex distinguishing coherent inputs from word lists, and to two right-hemisphere story effects in extended language regions. We also observed a context effect in the Slow Story condition, with neural responses remaining constant as the narrative unfolded while they increased in the SentenceList and WordList conditions. This effect was absent under Fast presentation, suggesting story-specific comprehension that is temporally constrained. Together, the findings identify temporal constraints as a key determinant of the neural signatures of narrative processing.

## INTRODUCTION

What are the neural mechanisms underlying the processing of meaning in large, complex contexts? A broad literature has investigated the neural bases of single-word processing and phrase-level composition [1–6], including their extensions to sentence and story comprehension [7,8], where information is integrated over longer timescales. At the sentence-level, the integration of word meanings elicits increasing activity in language selective regions over the course of a sentence, with highest amplitudes at the end [9–11]. If this finding reflects the general accumulation of meaning, we expect a similar pattern at the narrative level, with signals in language cortex increasing as a story progresses. Alternatively, previously reported increases could be a sentence level phenomenon, always resetting at the end of a sentence.

Most theories of discourse processing do not merely consider narratives as long sentences. Instead, they posit the continual creation and updating of ‘situation models’. Situation models are mental representations of the world that allow us to interpret events, whether they arise in a narrative, a larger discourse or just everyday experience [12–14]. Situation model construction extends beyond left-lateralized language cortex, engaging right hemisphere (RH) homologs [15] and the Default Mode Network (DMN) [16].

Many studies have contrasted narratives with smaller representational units such as sentences and individual words. In their seminal paper, Xu et al. [17] contrasted random words, shuffled sentences and intact stories, finding that fMRI activation became increasingly bilateral as representational complexity increased. This pattern has since been replicated in studies using narratives to explore the neural mechanisms of situation model processing [18], to understand language processing at different timescales [19] and to naturalistically engage semantic representations [20–22].

Few studies have capitalized on the fact that successful narrative comprehension also requires sufficient time to contrast and update situation models. Thus, a narrative should engage the brain more broadly than isolated sentences, and this deeper processing likely requires more processing time. Independently of the size of the representation, presentation speed could modulate depth of processing. In this study, we combined the two approaches, varying both the size of the representational units (words, sentences, stories) and the speed at which the stimuli were presented. While visual presentation speed has been varied in at least three prior studies [23–25], we do not know how presentation speed affects processing across multiple representational levels.

We also introduce an adaptation of the so-called rapid parallel visual presentation (RPVP) paradigm to the presentation of a natural narrative stimulus. In this paradigm, a multiword expression is presented at once, for a few hundred milliseconds [26]. The experience is highly ecologically valid in the modern digital world, since this type of at-a-glance perception is how we often read notifications on our phones, road signs while driving, ad slogans and even text overlays in social media videos. This approach contrasts with most prior visual sentence-comprehension work, which has used rapid serial visual presentation (RSVP), presenting one word at a time in a way that does not resemble natural reading. We built on a series of recent MEG studies using RPVP for the study of basic syntactic and semantic processing [27–30], taking advantage of the natural reading experience it offers both at slower and faster speeds.

To investigate how narrative processing is distinct from sentence and word-level processing, we collected magnetoencephalography (MEG) data while participants read two personal essays delivered in RPVP using 1-4 word chunks. The stimuli were presented at three levels of structure (intact story, sentence lists, and word lists) and two levels of speed (Slow at 800ms per chunk and Fast at 450ms per chunk), with the timing guided by prior electrophysiological literature on sentence processing. We first used mixed-effects ANOVAs to identify functional Regions of Interest (fROIs) sensitive to structure in the two speeds. To characterize the effects of incremental story context within the identified regions, we fit a generalized linear model with position within a block and condition as the main regressors, enabling a dissociation between neural signals related to story development from those related to time on task. Together, these analyses allowed us to delineate structure-related responses, their sensitivity to speed and their evolution across the unfolding narrative.

## MATERIALS AND METHODS

### Participants

35 participants (19 women, 16 men and 0 non-binary; mean age = 27 years) took part in the MEG experiment. They were recruited online using New York University’s SONA system. The sample included both paid adults compensated 15$/hr for their time and undergraduate students who received course credit. All participants were right-handed, highly proficient English speakers (mean proficiency ± SD, 0.9727 ± 0.0469) and had normal or corrected to normal vision. Two participants were excluded due to overall accuracy below 60% in our chunk recognition task. One participant reported having ADHD and another undiagnosed dyslexia.

### Design

As shown in Fig 1, the experiment followed a 3 x 2 factorial design with Structure (WordList, SentenceList, Story) and Speed (Slow, Fast) as main factors. Our protocol was designed to minimize top-down effects from prior exposure to the narrative while maintaining good lexical and sentence-level control. Accordingly, two distinct stories were used for the Slow and Fast presentations (counterbalanced across participants), and the three levels of structure were constructed from the same narrative. To prevent narrative-level information from influencing the processing of our scrambled stimuli, WordLists were always presented first, then SentenceLists, and finally the intact Stories. Furthermore, presenting the Slow rate before the Fast rate facilitates contextual integration in the Fast condition [25], so Fast presentations always preceded Slow presentations.

**Fig 1.**
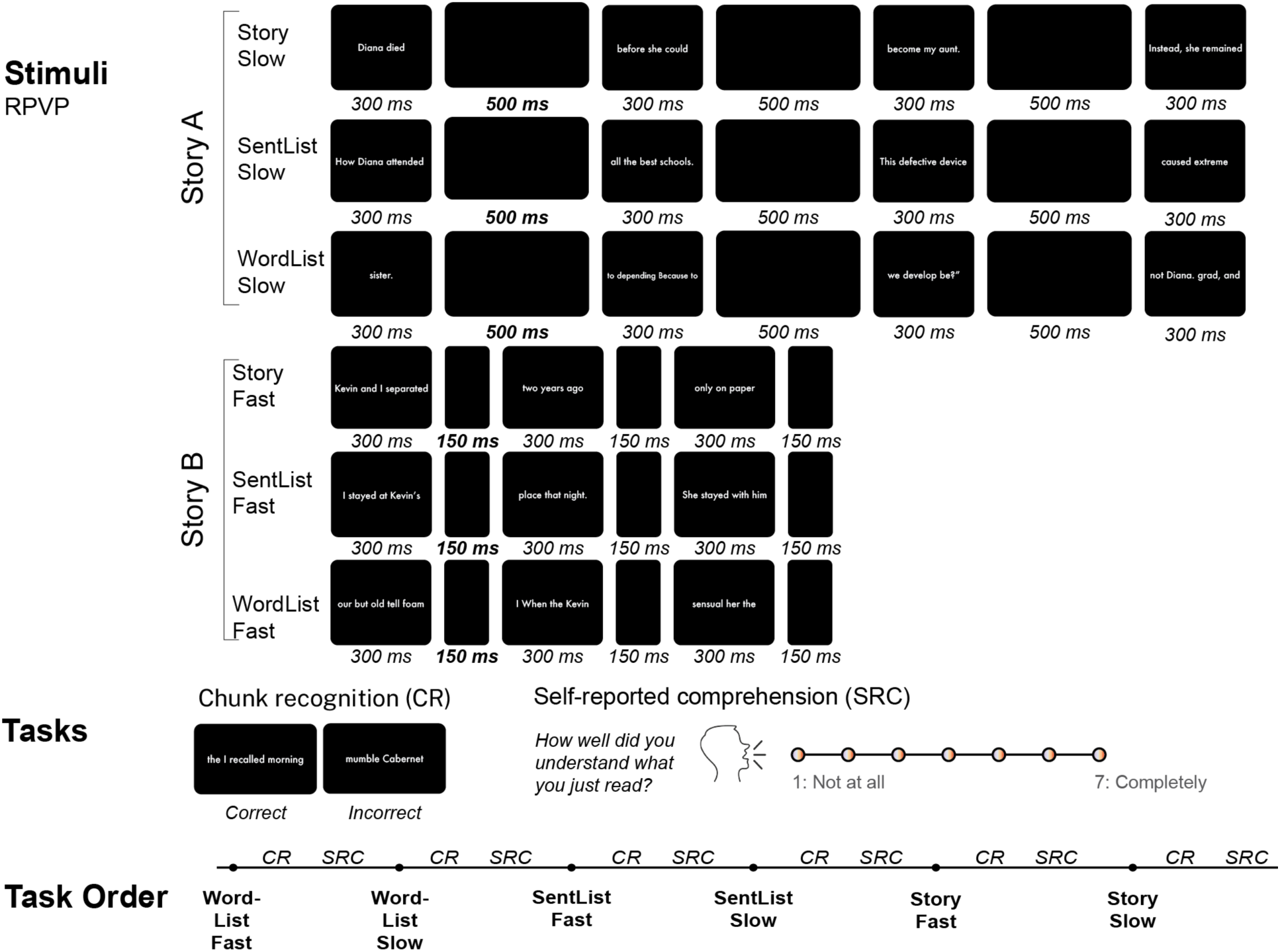
Stimuli, Tasks and Task Order. Our design had a 3 x 2 design with Structure (WordList, SentenceList, Story) and Speed as factors (Fast and Slow). We additionally had a chunk recognition task and a self-reported comprehension score.

There were 2987 trials in total. The number of trials per condition was equal to the number of chunks in each story: participants with Story A in the Slow condition had 537 trials in every Slow condition and 456 trials in every Fast condition. This was reversed with participants with the opposite order.

Stimuli were presented in 1- 4 word chunks using the Rapid Parallel Visual Presentation (RPVP) paradigm [26]. In the Slow condition, each chunk was onscreen for 300ms, followed by a blank screen of 500ms. This timing enables the detection of all traditional sentence processing stages up to the P600 [31] and replicates prior RPVP MEG experiments [27–30]. The Fast presentation represented a 56% temporal compression of the Slow condition. To be able to cleanly compare evoked activity across speeds, chunks were also onscreen for 300ms during the Fast condition. However, they were followed by a shorter 150ms blank screen. This timing allows us to minimally detect early MEG components such as visually evoked responses from 100-200ms [32] and early composition-related activity in the left temporal lobe. In studies presenting words serially, composition-related activity canonically peaks at 200-300ms [33], however, in RPVP, it has commenced as quickly as 130ms . After each block, participants rated their comprehension on a likert scale and completed a chunk recognition task (see Comprehension Tasks below).

### Stimuli

Our study employed two personal essays and a book chapter obtained from our collaborating author Alessandra Ranelli: “Two Kisses We Never Talked About” (TK), “My Mother’s Sister” (MMS), and “Jack Judd Ain’t Got No Love Songs” (JJ). TK was previously published in the New York Times Modern Love column. The other two stories were unpublished manuscripts. We chose these personal essays because of their naturalistic and conversational style. TK and MMS were used as the story stimuli while JJ chunks were used as filler items in the chunk recognition task (intended to elicit no-responses). TK was 456 chunks long and MMS was 537 chunks long. The stories were counterbalanced across participants which prevented differences in power across presentation speeds during group-level analyses.

All three stories were chunked into smaller phrases using Stanza [34] and a custom python script. The script chunked the text into its largest syntactic constituents while constraining the chunks to be less than 23 characters long (mean character per chunk ± SD, 16.90 ± 3.90; mean words per chunk ± SD, 3.40 ± 1.03). The chunks were then manually edited for flow and readability to make the reading experience as pleasant as possible. SentenceList versions of the texts were obtained by randomly shuffling the sentences of the story for each participant. To create the WordList versions, we sampled the original texts with replacement and reconstructed chunks with random words and punctuation. Additional constraints were that the new chunks contained around the same number of characters as the original chunks and were under the maximum character limit (23 characters). Less than 10% of words were repeated across the three stories.

To detect significant differences in chunk length, we ran an ANOVA on the number of words and the number of characters in each chunk using Story (MMS, TK) and scrambled (True, False) as factors. We included both the intact and word-scrambled versions of the stories we presented in the experiment. There was a small effect of scrambling for character length (F_1, 1986_ = 59.92, p < .001; 11_2_ (partial) = 0.03) and number of words (F_1, 1986_ = 30.78, p < .001; 11_2_ (partial) = 0.02). However, this only meant that the scrambled stimuli were 0.2 words (or 1.2 characters) longer than the intact versions. We also found a small effect of story (F_1, 1986_ = 38.26, p < .001; 11_2_ (partial) = 0.02) on the character level: TK was around 1.16 characters shorter than MMS. However, we did not find significant differences in word length between the two stories.

### Comprehension tasks

Measuring incremental online comprehension during naturalistic paradigms is difficult: asking detailed questions during the experiment interrupts participants’ experience and asking for continuous ratings requires that participants take a step back to reflect on their experience, creating a dual task scenario. We used two tasks to evaluate if our speed manipulation affected depth of processing and comprehension: a self-reported comprehension score and a chunk recognition task.

Recognition memory can be used as a surrogate measure of comprehension [25,35]. Our Chunk Recognition Task followed every experimental block: 15 chunks from the preceding story block and 15 chunks from the filler story (JJ) were presented to the participants. None of the chunks repeated across blocks. Chunks included in the task following the WordList conditions were also scrambled. The prompt (“Please indicate if the following chunks were included in the previous text.”) was onscreen while participants reported if they recognized the chunks from the previous block. Responses were either Yes or No and were indicated using a button box in the MEG. Then, participants rated their comprehension out loud using a 7-point Likert scale (Prompt: “How well did you understand what you just read?”). We validated the chunk recognition measure by correlating it with our participants’ self-reported comprehension.

### Procedure

After providing informed consent, each participant’s head shape and the positions of five marker coils and three anatomical landmarks (nasion and peri-auricular points) were digitized using a Polhemus FastSCAN system (Polhemus, Vermont, USA). The position of the participant’s head relative to the MEG sensors was determined via the position of the marker coils at the beginning and end of the MEG experiment.

Outside of the MEG chamber, participants were led through a practice session that consisted of a short 34-chunk story written by ChatGPT in the style of a Modern Love column story. The practice story was followed by two sample chunk recognition trials and the comprehension rating task. The practice story was presented at the Fast presentation speed to get participants used to the RPVP paradigm. Participants were encouraged to do the practice session as many times as needed before starting the MEG experiment.

The stimuli were presented with Psychopy in python [36] (version 2022.2.5). Each chunk was presented in white text on a grey screen in Futura font with a height of 7% of the presentation window at a mean horizontal visual angle of 4.784° (SD = 1.134). The screen was placed around 50cm away from the participants. Participants could take breaks before and after each reading block and were instructed to limit their eye movements while the text was flashing on the screen. The recording session lasted around 1h20min per participant.

### MEG data acquisition and preprocessing

Magnetoencephalography data was recorded using a 157-channel axial gradiometer whole-head MEG system (Kanazawa Institute of Technology, Kanazawa, Japan) at a sampling frequency of 1000hz with an online bandpass filter from 0.1hz to 200hz.

Participants lay down in a dimly lit magnetically shielded room. Triggers were sent with each chunk and a photodiode was used to estimate the delay between the onset of the chunks and the triggers. During preprocessing, trial onsets were corrected using this delay.

Once collected, the data were noise reduced in MEG160 (Yokogawa Electrical Corporation and Eagle Technology Corporation, Tokyo, Japan) with a Continuously Adjusted Least-Squares Method algorithm. The raw data were then preprocessed using MNE-Python 1.10 [37]. First, we filtered the raw data using a 1-40hz bandpass IIR filter; the 1Hz high pass filter is typically necessary given Manhattan noise levels. We then used Independent Component Analysis to identify and remove known MEG artifacts such as blinks, heartbeat, as well as radio noise and other environmental noise. Trials were then segmented into epochs spanning -100ms to 800ms relative to stimulus onset for the Slow presentation and 0 to 450ms for the Fast presentation as we initially wanted to use a prestimulus baseline for baseline correction. Since the prestimulus period cannot be analyzed during the Fast presentation, we started our analyses at 0ms and demeaned our trials across the whole epoch. The Fast and Slow presentation conditions were separated and demeaned independently. Trials with a peak-to-peak amplitude exceeding 3000 fT after noise reduction were automatically rejected and any bad channels were removed (Max = 9, Min = 1, Mean = 3.3).

### Source reconstruction

For participants without a structural MRI, the FreeSurfer [38] average brain was scaled to fit their digitized headshapes and fiducials. We used the symmetrical version of Fsaverage (Fsaverage-Sym) to be able to detect hemispheric differences. MEG data were coregistered to the participant’s scaled (N = 26) or anatomical MRI (N = 7) using the anatomical landmarks. We then computed a source space with 2562 vertices per hemisphere. Forward solutions were computed for each participant using a Boundary Element Model. A channel noise-covariance matrix was calculated using two minutes of continuous resting state data that underwent the same ICA, filtering and bad channel removal pipeline as the rest of the experiment. We estimated an inverse solution from the forward solution and covariance matrix using an SNR of 3 for ANOVA analyses of Structure and Speed and an SNR of 2 for the single-trial GLM analyses addressing story progression.

We projected both single-trial sensor data and condition-averaged evoked responses into source space, using the respective inverse operator, to obtain L2 minimum norm source estimates. This yielded noise-normalized Dynamic Statistical Parameter Maps (dSPM).

### Behavioral data analyses

#### Chunk recognition task

Trials with response times shorter than 200ms or longer than 8000ms were excluded from analysis. We additionally removed any trials with response times more than 3 SD away from each participant’s mean response time.

To determine the effects of Speed and Structure, we fit two generalized linear mixed effect models (GLMM): a model with a binomial distribution that predicted accuracy with Structure and Speed (formula: accuracy ∼ Structure * Speed) and another that predicted the log-transformed reaction times with our Structure and Speed factors. Both models included subjects and trials as random effects and the order in which they saw the stories (group) as a fixed effect.

#### Self-reported comprehension

Self-reported comprehension scores were z-scored across conditions for each participant before statistical analysis. We then fit a linear mixed effects model with the same structure as the one we ran on our chunk recognition data to predict our z-scored comprehension ratings (formula: comprehension ∼ Structure * Speed + (1 | Participant)). We used Chi-square tests (ξ_2_) for the logistic regression model and F-tests for the general linear models to assess our models’ significance.

### MEG data analyses

#### Effect of Structure for the two Speeds

Our first goal was to identify Structure-related effects in the Slow condition and then evaluate their replication under Fast presentation. Based on established fMRI findings showing increased neural activity for structured narratives relative to scrambled word lists or sentences, we expected greater structural coherence would be associated with stronger neural responses.

We ran two 3 x 1 mixed measures ANOVAs with Structure (WordList, SentenceList, Story) as a fixed factor and Subject as a random factor. We used the whole brain as a search area. The ANOVA was conducted at each vertex and timepoint from 0-800ms for the Slow condition and 0-450ms for the Fast condition. We established timepoints and vertices where there was a significant Structure effect through a spatiotemporal cluster-based permutation test [39] on uncorrected clusters of the resulting F-maps. Clusters had to have an F-value corresponding to an uncorrected p-value of 0.01, a minimum duration of 40ms and include at least 40 adjacent sources to be included in the analysis. Corrected cluster *p* values were estimated using 10,000 permutations and additionally FDR corrected across tests (n = 2). We ran follow-up post-hoc t-tests to establish significant differences between conditions and corrected for multiple comparisons using holm correction.

#### GLM analysis

Our ANOVA identified where stories were processed differently from sentences or random words, but collapsed over all chunks within a condition. Crucially stories, unlike sentences, have a structure that evolves over longer timescales. In fact, it is well-established that accumulating context over the course of both stories and sentences increases neural responses [9,10,40] such that there is higher activity at the end of a story or sentence than at the beginning. In order to evaluate how narrative processing changes over the course of a story, we ran a two-stage single-trial regression analysis [41] on our design.

We used trial number (Trial#) as our proxy for block progression, as trial number continuously increased within blocks but only in the Story conditions did it additionally track narrative progression. If narrative progression is reflected through a gradual increase in activity, then the effect of Trial# should be larger for the Story conditions relative to both the WordList and SentenceList conditions. Thus, our analysis focused on the interaction between Condition and Trial#.

Trial# was defined relative to the start of each block, such that the first trial in each condition was Trial0. The regressor was then scaled relative to the total number of trials in the block and centered before being entered into the analysis. We used Story as our reference level for the Condition regressor in the model.

Data were downsampled to 500hz to speed up computations. In the first stage, we fit an ordinary least squares regression model that predicted the dSPM activity at each vertex and each timepoint across trials (see Formula 1). Our analysis time window was the same as for our ANOVA (0-800ms) and we used the ANOVA results in the Slow presentation as a functional ROI. We selected significant clusters from both hemispheres and mirrored them onto the opposite hemisphere to get a balanced search area. For the Fast conditions, the analysis time window was shortened to 0-450ms. A total of two models were run per participant: one for each speed. The first stage resulted in spatio-temporal maps of β coefficients for each regressor.

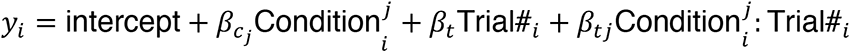

**Formula 1**: Formula for regression model.

In the second stage of analysis, we ran a one-sample spatio-temporal t-test to determine group-level effects of each regressor. Clusters had to have an uncorrected *p* value of 0.05, include at least 20 adjacent sources and last longer than 20ms to be included in the clustering analysis. Corrected cluster *p* values were estimated using 10,000 permutations and additionally FDR corrected across tests (n = 12). Both the GLM analysis and ANOVA were conducted with Eelbrain [42] (version 0.40).

## RESULTS

### Behavioral results

We presented scrambled and intact narratives at Slow and Fast speeds, to study the temporal constraints around narrative processing. Our participants completed a chunk recognition task and rated their comprehension of the stimuli, which we used to measure comprehension after each block. Chunk recognition accuracy was quite low (mean ± SD, 0.706 ±0.059) due to the difficulty of the task. Task accuracy was significantly correlated with self-reported comprehension across participants (*r* = 0.52, *p* < .001) indicating that our chunk recognition task successfully measured comprehension. An overview of the behavioral results across conditions has been provided in Table 1. Additionally, which story participants read in the Slow or Fast conditions did not affect their task performance.

**Table 1.** Average behavioral performance per condition for MEG sample (n = 33).

| Condition | Speed | Accuracy (au, M ± SD) | Comprehension (z, M ± SD) | Reaction time (ms, M ± SD) |
| --- | --- | --- | --- | --- |
| Story | Fast | 0.7505 (0.0862) | 0.8144 (0.3115) | 1808.785 (898.2514) |
|  | Slow | 0.7758 (0.0818) | 1.0511 (0.2566) | 1748.461 (905.7858) |
| SentenceList | Fast | 0.7061 (0.0775) | 0.1335 (0.3807) | 1783.319 (867.9767) |
|  | Slow | 0.7303 (0.1100) | 0.2554 (0.3283) | 1654.197 (834.0180) |
| WordList | Fast | 0.6545 (0.0740) | 1.0450 (0.3520) | 2417.250 (1171.4793) |
|  | Slow | 0.6192 (0.1096) | 1.2095 (0.2714) | 2064.920 (1044.8186) |

Structure significantly affected Chunk Recognition Accuracy ( $\chi^2(2) = 79.19, p < .001$ ), Reaction Time ( $F_{2, 5533.8} = 213.38, p < .001$ ) and Self-Reported Comprehension ( $F_{2, 191} = 719.00, p < .001$ ). Speed affected Chunk Recognition Reaction Time ( $F_{1, 5532.7} = 41.449, p < .001$ ) and Self-Reported Comprehension ( $F_{2, 191} = 719.00, p < .001$ ).

As predicted, Structure significantly affected behavioral performance (see Fig 2): participants had higher accuracy, self-reported comprehension scores and faster reaction time in conditions with higher semantic coherence. Further pairwise testing confirmed a graded difference across levels of Structure (*p* < 0.001). Specifically, Stories were better understood than SentenceLists, which in turn were better understood than WordLists. Although coherence decreased response time, there was no additional speed advantage for Stories over SentenceLists, suggesting a ceiling effect.

**Fig 2.**
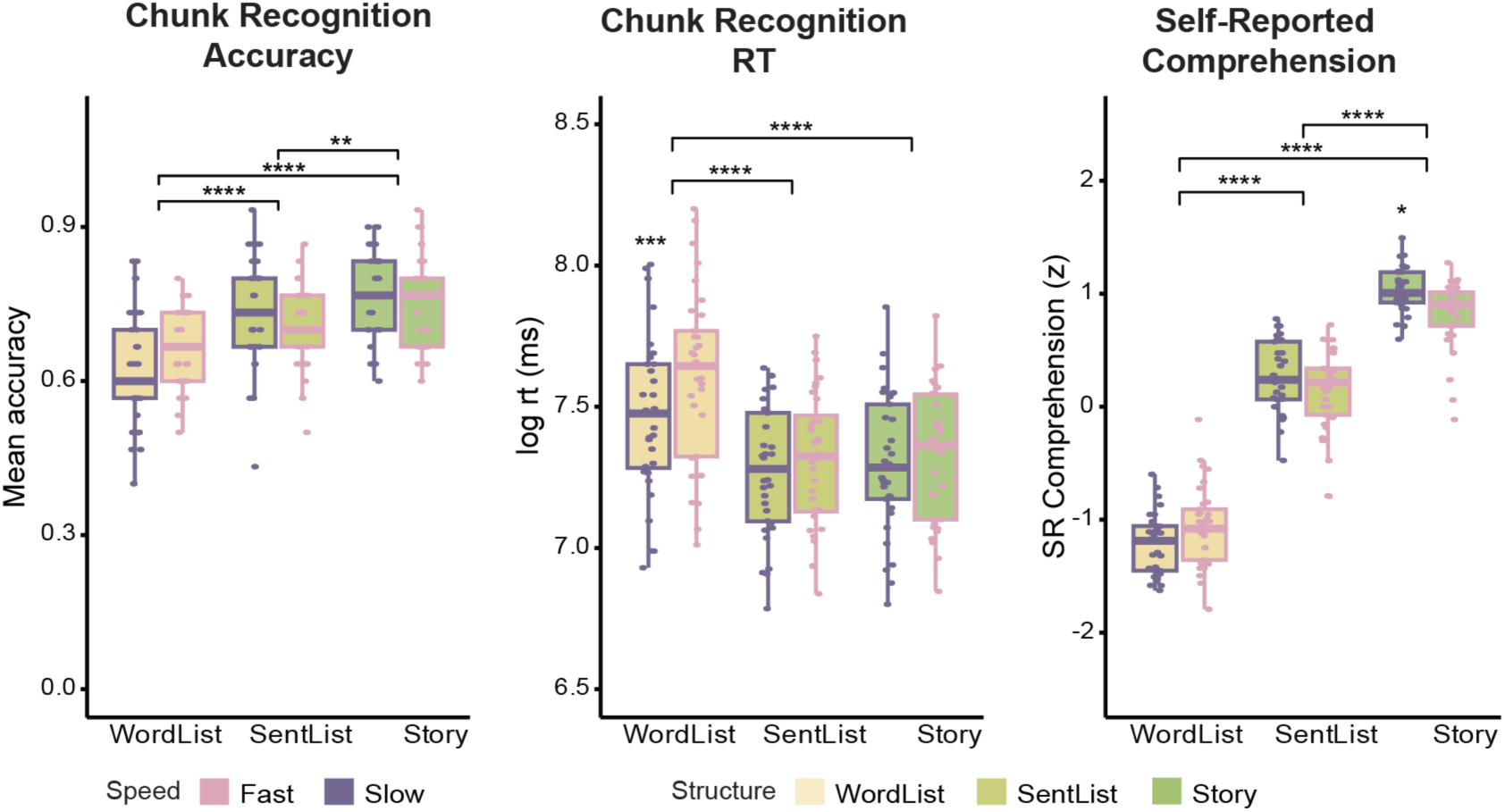
Behavioral results for MEG experiment (n=33). Comprehension scores and Task accuracy were higher in the Story condition than in the SentenceList condition (Accuracy: *z* = -3.32, *p* = .003; Comprehension: *t* = -13.33, *p* < .001), in the Story compared to the WordList condition (Accuracy: *z* = -8.75, *p* < .001; Comprehension: *t* = -37.41, *p* < .001) and in the SentenceList compared to the WordList condition (Accuracy: *z* = -5.47, *p* < .001, Comprehension: *t* = -24.08, *p* < .001). Reaction times were faster in both the Story (*t* = 17.314, *p* < .001) and SentenceList (*t* = 18.462, *p* < .001) conditions vs. the WordList condition. Significant differences between Fast and Slow presentation within one level of structure is indicated by a bolded asterix above the Slow condition. Asterix legend: . < 0.1, * p < 0.05, ** p < 0.01, *** p < 0.001, **** p < 0.0001.

While increasing presentation speed significantly increased response times (*t* = - 6.438, *p* < .001), both task accuracy and self-reported ratings were unaffected.

Significant interactions revealed that the effect of Speed was dependent on Structure: Fast presentation selectively increased response latencies for WordLists (*t*_5534_ = -7.132) and impaired comprehension for Stories (*t*_160_ = 2.942, *p* = 0.0427). Behaviorally, increasing presentation speed unveiled a diverging pattern of results: participants had slower response times to incoherent stimuli but reported understanding the stories better when given more time to process them.

### MEG results

#### ANOVA results

We conducted two mixed measures ANOVAs across the whole brain to identify where and when Structure effects emerged in the Slow and Fast conditions. We analyzed the Slow and Fast data separately for two reasons: first, epochs in the Slow and Fast conditions had different lengths. Second, we sought to identify Structure effects during maximal comprehension (Slow presentation), and determine whether these patterns persisted under increased temporal pressure.

Our results discussion will be focusing on two different types of effects: (i) **Structure effects,** a graded increase in activity across our three levels of Structure (Story > Sentence > Word) (ii) **Story effects**, a preferential increase in activity for the Story condition with no significant difference between Sentences and WordLists. Fig 3 is a comparative summary of the timeline and spatial extent of our results across Slow and Fast presentations, while Figs 4 and 5 contain our results per hemisphere.

**Fig 3.**
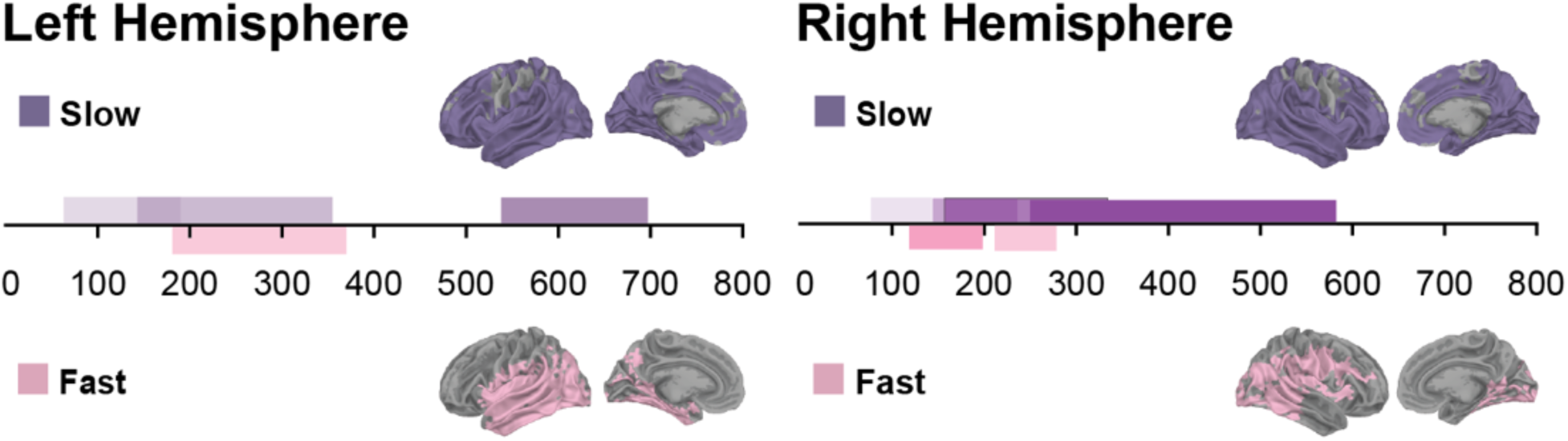
Comparative summary of timecourse and spatial extent of significant clusters across Slow and Fast presentation modes. Slow results (in purple) have been placed above Fast results (in pink) on the x-axis.

**Fig 4:**
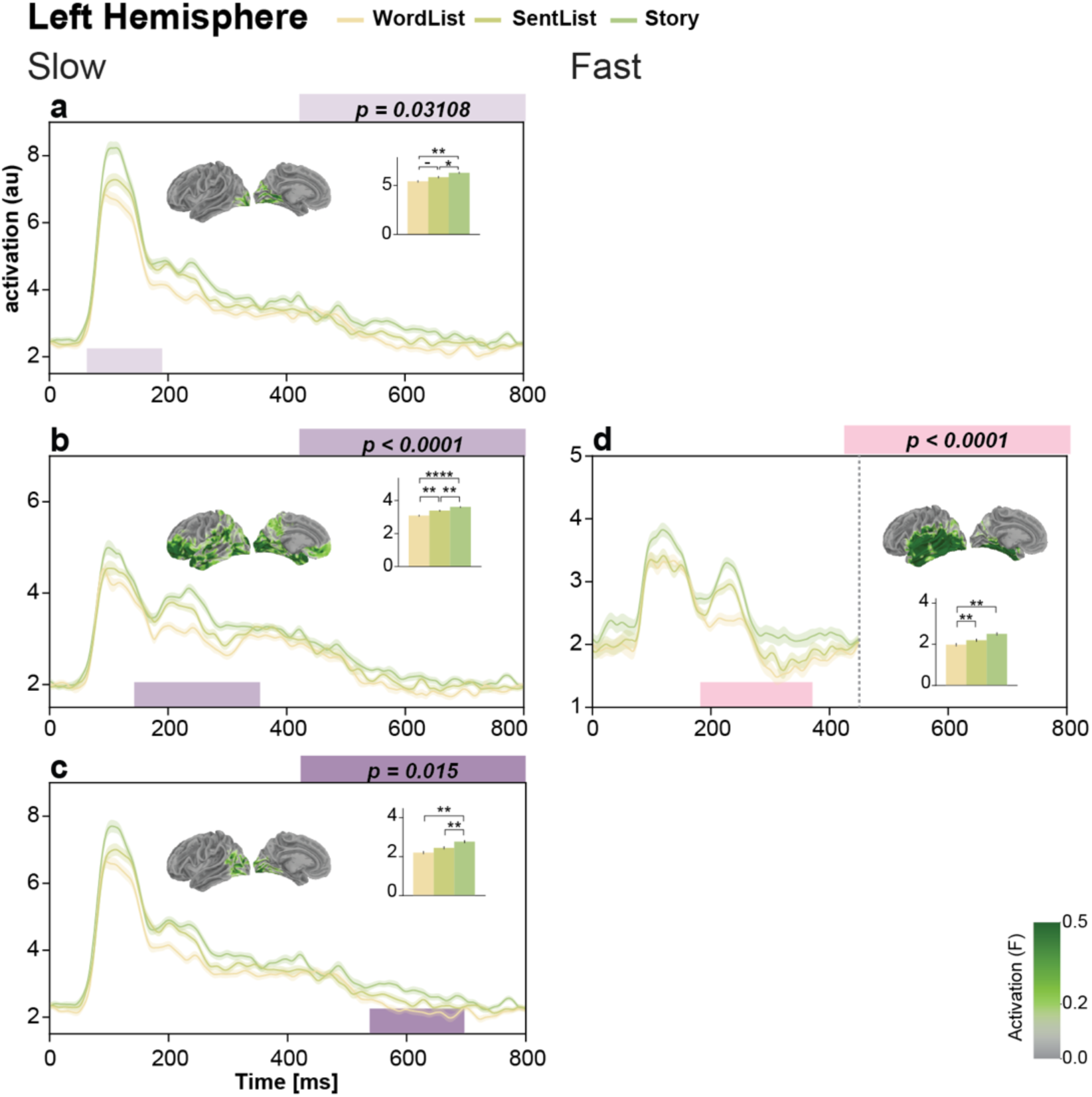
Left hemisphere results for Slow and Fast conditions. Fast results (in pink) have been placed to the right of their Slow presentation equivalent (in purple) where possible. a-d: Individual timecourses, mean amplitude across conditions and spatial extent of significant clusters. Asterix legend: . < 0.1, * p < 0.05, ** p < 0.01, *** p < 0.001, **** p < 0.0001.

**Fig 5.**
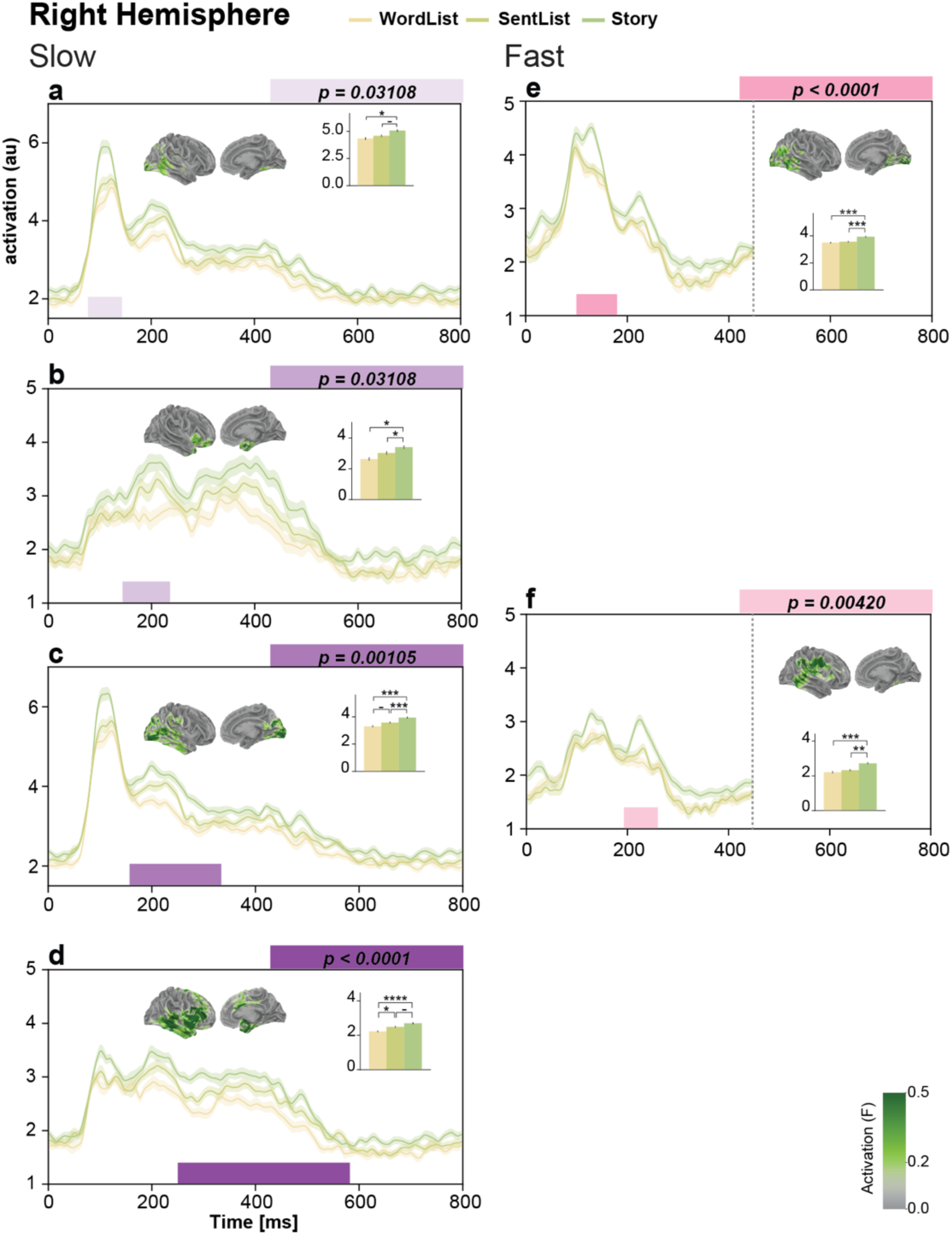
Right hemisphere results for Slow and Fast conditions. Results for Fast presentation (in pink) have been placed to the right of their Slow presentation equivalent (in purple) where possible. a-e: Individual timecourses, mean amplitude across conditions and spatial extent of significant clusters. Asterix legend: . < 0.1, * p < 0.05, ** p < 0.01, *** p < 0.001, **** p < 0.0001.

##### Slow condition

Starting with the Slow condition, in the left hemisphere, we found a large cluster (*p* < 0.0001; Fig 4a) showing a widespread Structure effect from 143 to 355ms that included dorsolateral prefrontal cortex, core language regions like the anterior temporal lobe or inferior frontal gyrus and as well as extended language regions like extrastriate cortex, ventromedial prefrontal cortex and precuneus. This cluster showed a threeway contrast between Stories, SentenceLists and WordLists such that more semantically coherent stimuli elicited higher activity.

Primary visual and visual association cortex exhibited early and late clusters suggesting both feedforward and feedback processing within the visual hierarchy. Both clusters showed a Story effect where SentenceLists and WordLists patterned together. The early cluster (*p* = 0.031; Fig 4a) was concentrated in V1 and lasted from 63ms to 190ms. The late cluster (*p* = 0.015; Fig 4c) lasted from 538ms to 697ms and included more of visual association cortex.

In the right hemisphere, we obtained a spatio-temporal signal progression from visual and posterior temporal areas through anterior temporal and frontal language regions until prefrontal cortex. Within this, the earliest cluster occurred at 77-143ms in the occipital pole and posterior temporal lobe (*p* = 0.03108; Fig 5a), with a significant difference only between the Story and WordList conditions, and a trending difference between the Story and SentenceList conditions.

At around 150ms, we observed a cluster that included most of the temporal lobe, visual association cortex and angular gyrus (157-334ms, *p* = 0.001; Fig 5c). This cluster showed a Story effect with a trending difference between SentenceLists and WordLists.

Finally, our longest cluster ranged from 250ms to 582ms in canonical language regions like the inferior frontal gyrus and middle temporal lobe (*p* < 0.0001; Fig 5d). This cluster differentiated between conditions with composition (Story, SentenceList) and the WordList condition with a trending difference between the SentenceList and Story conditions. We additionally found a Story effect in the temporal pole and inferior frontal regions at 144 - 236ms (*p* = 0.03108; Fig 5b).

##### Fast condition

We then evaluated the robustness of our Structure manipulation at the faster presentation speed by running the same 3 x 1 ANOVA in the Fast conditions. In the left hemisphere (LH), speed eliminated story-specific processing. Both our early and late Story effects in visual cortex were eliminated. Our Fast cluster (182 to 371ms, *p* < 0.0001; Fig 4d) resembled the broad language network response we found in the Slow condition (Fig 4b), including most of the temporal lobe as well as visual cortex but excluding frontal regions. This cluster had a later onset than its Slow condition equivalent and only distinguished between coherent (Story and SentenceList) and incoherent (WordList) conditions.

In the right hemisphere, we observed the same spatio-temporal signal progression as in the Slow condition where early effects in visual areas develop into later clusters in anterior language regions. The earliest cluster at 100-180ms (*p* < 0.0001; Fig 5e) resembled the first visuo-temporal cluster we found during Slow presentation with crucial differences. The Fast presentation cluster exhibited a visually stronger Story effect, which started later, and included more visual association cortex than its Slow presentation equivalent.

Our second Story effect during Fast presentation occurred at 193-260ms in middle temporal, inferior frontal and somatosensory areas (*p* = 0.004; Fig 5f) and did not resemble any cluster obtained for the Slow condition. We did not replicate the early temporal pole Story effect (Fig 5b) or the large language network response (Fig 5d) we found in the Slow condition.

### GLM results

Activation is known to be higher during later portions of a story compared to the beginning [17,40] and accumulating story context increases right hemisphere engagement more than the left. Our first round of analyses used evoked responses and so were unable to capture the temporal dynamics of narrative construction. To account for the cumulative effects of story progression on neural activity, we conducted a single-trial regression analysis. First, we formed an fROI from our significant clusters in the Slow condition. Our regression model used trial number (Trial#) as a proxy for block progression. Though each block had the same number of chunks across conditions, in the Story condition, block progression also tracked story progression. We expected this compound Block-Story effect to influence neural activity, so we used the Story condition as a baseline and focused our analysis on the interaction between Trial# and Condition. We then evaluated the group-level effects of our regressors within our fROI with a one-sample t test.

#### Main effects: WordList, SentenceList, Trial#

As expected from our ANOVA results, we found a main effect of both WordList and SentenceList across presentation speeds: fROI amplitude was lower in the WordList and SentenceList conditions compared to the Story condition (Fig. 6A). For the Slow presentation, the SentenceList decrease was concentrated in bilateral visual and temporal regions for the entire time window (LH: 0-800ms, *p* = 0.0138; RH: 0-800ms, *p* = 0.0210). In the Fast condition, the difference between SentenceList and Story was right-lateralized and strongly observed in inferior temporal as well as inferior frontal and occipital regions (RH: 0-450ms, *p* = 0.02998) with a trending left hemisphere cluster.

**Fig 6.**
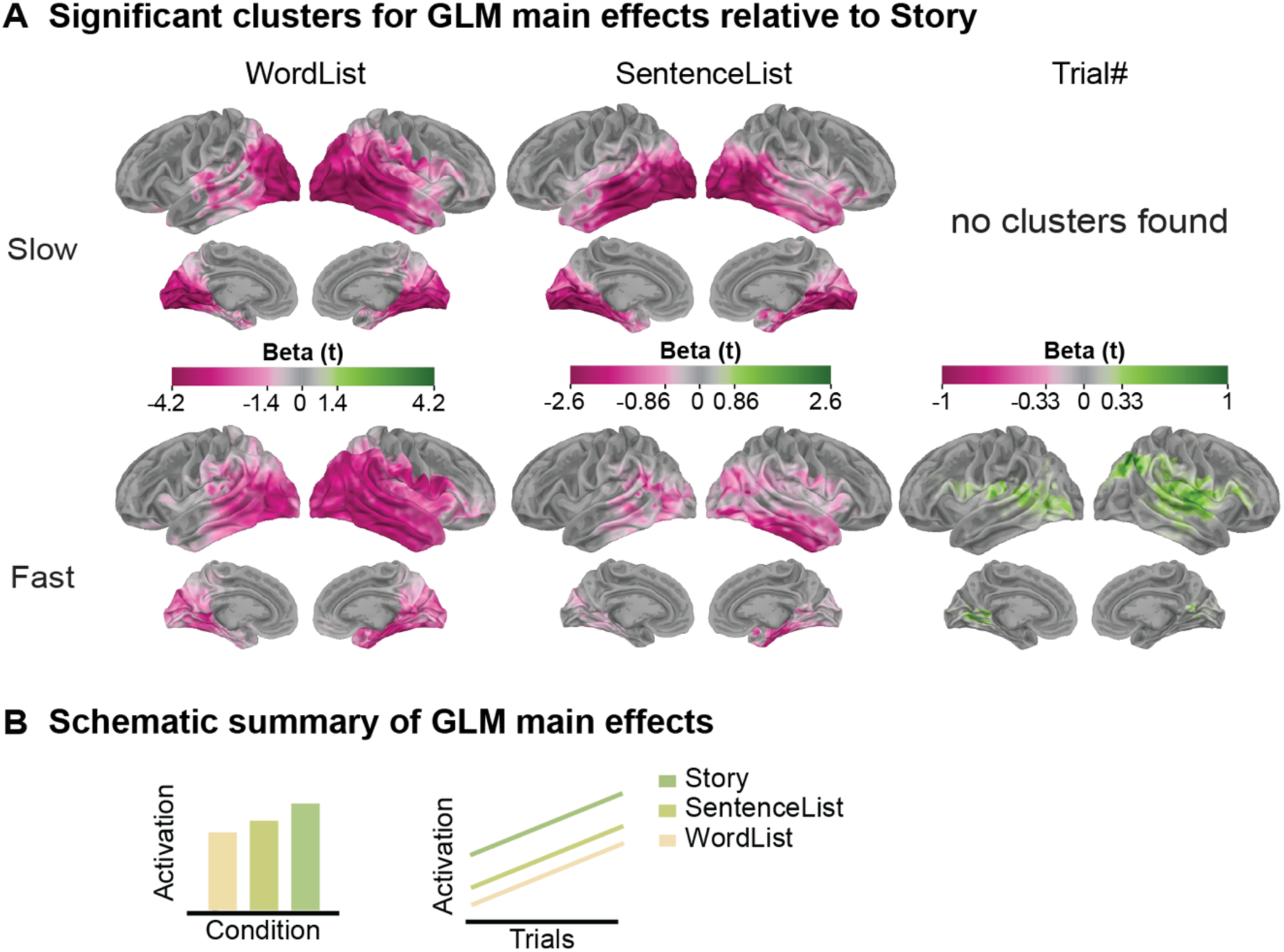
GLM model main effects. Trial# indexes block progression in all conditions but additionally tracks narrative progression only in the Story condition, which serves as the reference level in the model. A: Spatial summaries of β t-maps for each regressor.

Our WordList effect (β_max_(t) = 5.9) was larger than the SentenceList effect (β_max_(t) = 2.9). The WordList activity decrease was strongest in bilateral occipital cortex and right inferior frontal and superior parietal regions. During the Slow presentation, we observed two clusters that spanned most of our fROI excluding the superior frontal lobe and the left temporal pole (LH: 0-800ms, *p* = 0.0067; RH: 0-800ms, *p* = 0.0006). These same clusters were replicated in the Fast condition (LH: 0-800ms, *p* = 0.0067; RH: 0-800ms, *p* < 0.0001), with the additionaly contribution of the left posterior temporal lobe as well as most of the right ventral prefrontal cortex additionally contributed to the effect.

Effects for WordList and SentenceList are shown relative to the Story baseline (model Intercept); the Trial# regressor tests for a linear effect within the Story condition, for which no significant clusters were found in the Slow presentation, suggesting that narrative processing stabilizes responses and offsets a general increase over trials, whereas under Fast presentation this stabilization is reduced and Trial# effects emerge. B: Schematic summaries of main effects of Condition and Trial#. Trial# was only significant under Fast presentation.

We defined Trial# as the incremental effect of trial within each block. With Story as the reference level, the Trial# coefficient reflects the effect of trial number within the Story condition. A positive coefficient would indicate that neural activity in the Story condition was positively correlated with trial number such that later trials had higher activity than earlier ones. Reversely, a negative coefficient indicates that earlier trials had higher activity than later ones. We did not find a significant effect of Trial# in the Slow condition, contrary to the prediction of language processing models in which story progression elicits a steady increase in activation across trials. However, we did find a bilateral positive effect of Trial# in the Fast condition, replicating the increase in neural activity at later trials that we expected based on the literature. Our left hemisphere cluster was from 96ms to 298ms (*p* = 0.01748) and spanned left posterior temporal regions, the inferior frontal gyrus, the angular gyrus and visual association cortex. While our right hemisphere cluster had a similar latency from 114ms to 362ms, the effect was concentrated in middle temporal and inferior parietal regions (*p* = 0.01636).

#### Interactions between Trial# and Condition

We expected Trial# to have a stronger effect in the Story condition, where it additionally tracks the accumulation of narrative information. As such, the crux of our analysis was the interaction between Trial# and Condition. For the Slow presentation, we found that both WordList and SentenceList had a significant interaction with Trial# (Fig 7A). The WordList:Trial# interaction yielded 4 homologous clusters across both hemispheres. We observed two early clusters from ∼0-330ms (LH: 0-334ms, *p* = 0. 0.0067; RH: 0-342ms, *p* = 0.00998) and two late clusters from ∼330-800ms (LH: 338-800ms, *p* = 0.0064; RH: 332-800ms, *p* = 0.00641 which all spanned the entire fROI. The interaction between SentenceList and Trial# yielded one early left-lateralized cluster from 14ms-316ms (*p* = 0.02224). The effect was spatially similar to the WordList:Trial# effect though sparser in lateral prefrontal cortex.

**Fig 7.**
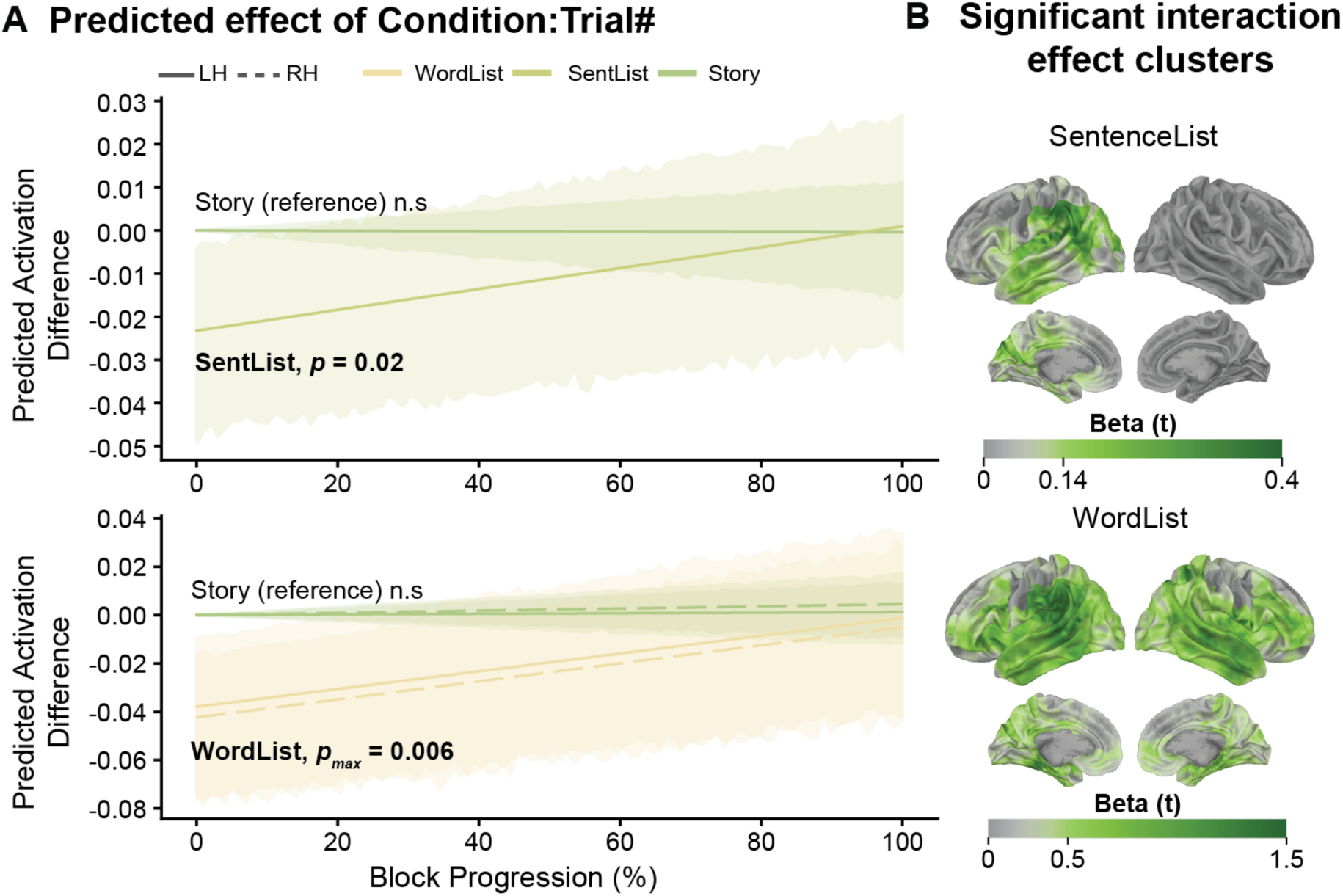
GLM interaction results showing increases in SentenceList and WordList conditions but not in Story. A: Predicted cluster amplitude for the Condition:Trial# interactions, averaged over significant vertices and timepoint for each cluster. 95% confidence intervals across subjects are indicated by a shaded bar. Interactions effects are indexed by changes in SentenceList and WordList relative to the Story baseline and the Story condition is plotted for reference. The same vertices and timepoints used to plot the conditions of interest (WordList, SentList) were selected to index the effect of Trial# in the Story condition. The significance of each interaction has been placed next to the condition label. If a condition had multiple clusters, the largest p-value has been indicated. B: Spatial summary of WordList:Trial# and SentenceList:Trial# effects.

Our two Condition:Trial# interactions were observed in the absence of a main effect of Trial#, indicating that neural activity significantly increased across trials during the SentenceList and WordList conditions but remained relatively constant during the Story condition (Fig 7A). This pattern suggests that the steady increase in activation across trials reported in prior work is not specific to narrative processing and may instead reflect block-internal factors such as effort, fatigue or shifting processing demands rather than story-specific processing, with the additional context in the Story condition mitigating or offsetting this increase.

In contrast to Slow presentation, there were no significant interactions between Trial# and Condition during Fast presentation: Trial# was positively correlated with neural activation and influenced all conditions equally. Our results indicate that condition-level differences due to narrative content are eliminated when there is limited time to process the input, consistent with the hypothesis that a linear increase in activity across trials indexes block-specific effort rather than story processing.

## DISCUSSION

Today’s growing consumption of content at increasing speeds raises fundamental questions about the limits of language processing. How much can we really understand from a social media video played at 2x speed? While prior work has characterized the spatiotemporal dynamics of word and sentence-level processing, much less is known about how temporal pressure affects narrative comprehension. Our results provide evidence that speed selectively impairs behavioral performance while attenuating neural responses. We provide a detailed picture of narrative-comprehension dynamics, consistent with a facilitatory effect of incremental context while providing further evidence that narrative comprehension is not a non-linear process.

### Information flow across levels of structure

In the Slow presentation condition, we found a bilateral fronto-temporal network that was engaged across early visual to core lexico-semantic sentence processing stages (∼90-600ms). Notably, in the right hemisphere we saw a three stage progression along the temporal lobe from 90ms to 600ms. Activation progressed from visual and visual association areas to fronto-temporal language regions. This spatio-temporal progression was also accompanied by an evolving sensitivity to our levels of Structure. A preferential story response at 90ms developed into a sustained response to semantically coherent conditions (Story, SentenceList) over shuffled WordLists at 250ms.

Our results are in line with growing evidence of a principal connectivity gradient [43,44] opposing primary unimodal cortices on one end with complex multimodal association areas. This gradient presents a natural processing hypothesis where the flow of information from sensorimotor to transmodal areas represents a progression from basic perceptual to more abstract forms of processing. Similarly, work on temporal receptive windows [45,46] proposes a cortical hierarchy processing linguistic material of increasing size. Sensory areas have the shortest windows, tuned for quick processing of perceptual features. Transmodal areas like the DMN have longer, paragraph-long windows that integrate contextual information over longer timescales. Crucially, information is integrated in short-windowed sensory areas before progressing to long-windowed multimodal areas, as reflected through a connectivity lag across cortical networks [19].

Although this study was the first to use RPVP with narrative stimuli, previous MEG-RPVP studies had already identified two phases of structure-driven processing for short parallel 2-4 word sentences: an early phrase-structure-sensitive phase in left posterior temporal cortex, beginning as early as 130ms [27] followed by a complexity-sensitive second phase in which top-down information can engage repair processes [28,29].

Here, by contrast, structure effects emerged much earlier, at 70-80ms, and were driven by a strong bilateral response in visual and visual association cortex with a robust preference for narratives. A key difference from the earlier MEG-RPVP studies is that in both our structured conditions, sentences and stories, the relevant representational units extended beyond the presented text chunks: each chunk was typically part of a larger expression and therefore came with context. This broader contextual embedding is the most likely source of the very early structure sensitivity observed here.

The impact of context also helps explain why the Story condition elicited such a strong visual response even though identical chunks were used in the SentenceList condition. Growing evidence suggests that primary visual cortex is involved in forming contextually-enriched bottom-up representations of complex visual stimuli. In a rapid scene detection paradigm, the semantic complexity of a scene enhances reaction time [47]. Regarding language processing, visual and visual association cortex have been shown to correlate with a model grouping word-forms within and across sentence contexts [48] and the visual network contributes to predicting self-relevance during personal narrative reading [49]. The timing of our early visual effects in the left hemisphere suggests that both feedforward and feedback processes are at play, with visual cortex projecting contextualized representations to specialized language regions. This interpretation is further supported by evidence that visual association cortex has longer temporal receptive windows than V1, making higher order visual areas plausible sites for context integration [45].

### Effect of Speed on language processing and comprehension

Previous studies have used presentation rate to determine processing bottlenecks in the language system, most often by increasing the speech rate of spoken words or sentences [50–52]. Though comprehension proportionally decreases with speed, speech can remain intelligible even at a 50% compression rate. Fewer studies explore this effect in the visual domain, often using rapid serial presentation where one word is presented at a time. In this study, we combined a novel parallel presentation paradigm with hierarchical levels of structure to characterize temporal constraints on language processing at multiple representational levels.

In line with the existing literature [53], the amplitude of our responses across conditions was lower in the Fast than the Slow condition and the latency of our Structure effects was delayed by about 50ms. Previous ERP evidence supports this latency shift: the N400 peaks 80-100ms later when words are presented at 10 words per second than when they are presented at 1.4 words per second [54].

#### Hemispheric differences in the effect of presentation speed

Faster presentation eliminated story sensitivity in the left hemisphere, indicating that the left hemisphere was especially vulnerable to temporal compression. This pattern is consistent with prior ERP evidence that faster presentation prevents semantic facilitation during the integration of words into a larger context. Wlotko and Federmeier [25] showed that increasing presentation speed weakens the N400 effect of implausible words that are semantically related to predicted words.

In the right hemisphere, story-specific clusters were more spatially restricted under Fast compared to Slow presentation. Activity was concentrated in middle temporal and angular gyrus regions and most robust to speed in posterior temporal cortex. The right hemisphere’s contribution to language processing remains debated. Some models view it as simply mirroring LH activity, whereas others propose a more specialized role in situation model construction [55] and discourse representations [56,57]. Relatedly, studies of discourse-level features such as coherence or inference generation have reported homologous activity across hemispheres [58]. Our results lean toward the specialization view, since story-specific responses were more robust to speed in the right hemisphere while disappearing in the left.

Across both hemispheres, Fast presentation completely eliminated dorsolateral prefrontal cortex (dlPFC) activation. Lerner et al. [46] varied the rate of auditory stories from 50% to 200% compression and found decreasing reliability in neural responses across subjects as speed increased; notably, no significant intersubject correlation was observed in medial or dorso-lateral prefrontal cortex when the story was presented at twice normal speed. The dlPFC has been linked to coherence processing and inference generation [58]. Coherence monitoring ensures that upcoming sentences are in line with the current situation model and is thought to be a default component of discourse comprehension. Slower presentation rates may therefore make coherence monitoring easier, especially for information unfolding on longer timescales. Consistent with a temporal-limitation account, Baumgarten and colleagues [59] found that while many regions show information bottlenecks, frontal sensors have temporal bottlenecks due to fixed-length integration windows: with less time available, these windows integrate less information.

Behaviorally, our results pose an interesting problem. While we observed a strong effect of speed on neural responses, it did not translate into an equally clear behavioral effect. Our participants’ responses slowed down during Fast presentation, but only when the content was not coherent, that is, in the WordList condition. Nevertheless, during the post-experiment debrief, participants reported struggling to understand the stimuli at faster speeds. One possible reason for this discrepancy is that our chunk recognition task may have tapped different memory processes depending on the stimulus type. Dual process models of recognition [60,61] distinguish between surface-level familiarity from deeper contextual recollection. Recognition memory is a fast-acting process that relies on the perceptual features of an object, while recollection memory is slower and can be facilitated by semantic coherence and contextual information.

Participants could have been relying on perceptual judgments in the WordList condition, which became much harder at a faster speed. However, a purely perceptual encoding of the SentenceList and Story conditions is much harder. The added context encourages a deeper level of processing which in turn affects recognition accuracy [62]. Since the amount of contextual information is the same across speeds within each level of structure, this may explain why speed did not affect accuracy in the structured conditions. This interpretation is further supported by the reaction time ceiling effect between SentenceLists and Stories, even though the depth of processing likely differed between those conditions. Overall, speed appears to reduce story-specific processing and attenuate neural response amplitudes in ways that are better reflected in subjective experience than in overt task performance.

### Is narrative comprehension a linear process?

To capture the various structure building processes that occur during narrative comprehension, such as situation model construction or the incremental updating of narrative representations, we conducted a GLM analysis that compared internal block progression across conditions, using the Story condition as a baseline. We used block-specific trial number (Trial#) as a proxy for block progression and tested for differences in the influence of Trial# across our conditions (Condition:Trial#). We found that Trial# was associated with a linear increase in the amplitude of neural responses across our fROI within the first 300ms of each trial across presentation speeds. This effect was specific to the WordList and SentenceList conditions during Slow presentation. Under Fast presentation, Trial# had the same positive effect across all conditions.

Previous evidence supports the idea that increasing constituent size linearly drives an increase in neural responses [11,63] and that narrative construction leads to an increase in activation across sentences and paragraphs [19]. In contrast, we found that increasing neural activity across a block depends on both block content (Condition) and presentation speed. It does not track story progression. The effect of Trial# was largest in regions that are typically associated with language processing in general, as opposed to narrative processing specifically, such as the posterior temporal lobe and the inferior frontal gyrus. According to Event Segmentation Theory, event segmentation is a constant, online process which additionally serves as a form of cognitive control, allowing for better resource allocation [64]. During narrative comprehension, event segmentation results in increased activity in posterior medial and prefrontal cortex around event boundaries [65]. Additionally, participants are highly sensitive to both the causal [66] and temporal structure of events within a narrative. Thus, comprehension is better characterized as a dynamic succession of cognitive state changes [67] which balances both information accumulation and contextual integration. Our results support a view of story comprehension where it is not a simple linear activity increase with each new piece of information. Viewing linguistic input as information or context accumulation fails to consider the natural event-related dynamics that dictate our perception and influence our comprehension.

Existing neuroimaging work on narrative processing has implicated the right hemisphere for the processing of discourse-level representations. However, our Trial#-related effects were more bilateral [17,40] and widespread than previous findings and both Condition:Trial# interactions occurred within the first 300ms of the trial. During this initial, possibly bottom-up stage, the presence of contextual information in the Story condition may facilitate processing and offset a potential block-fatigue effect.

Finally, the lack of interactions under Fast presentation suggests that these condition-related differences are dependent on having enough processing time to compute the input. The idea that constraints related to internal capacity limits, lead to an increase in activation or effort is not novel. Our results are consistent with the idea that language processing becomes more demanding with larger units, as proposed by capacity theories of language processing [68]. However, event segmentation naturally reduces stories into smaller units, facilitating their processing. At increasing speeds, the contextual story effect is reduced, such that there is still some degree of preferential story processing, but the depth with which the story is processed is different. While our behavioral results do not show a uniform effect of speed, neurally, we observe a reduction in overall tracking of Structure.

## CONCLUSION

Our goal was to characterize how linguistic structure and presentation speed shape the neural dynamics of narrative comprehension. We found that behavioral performance, subjective experience, and neural responses do not align in a straightforward way during narrative comprehension. Our results showed a spatiotemporal progression of activity along the temporal lobe that mapped onto the complexity of information being represented. Increasing presentation speed attenuated story-specific neural responses and had differential effects across hemispheres, eliminating story-specificity in the left hemisphere while preserving more restricted effects in the right. Finally, our results do not support a simple account in which neural activity increases linearly with accumulating information; instead, activity across trials depended on condition and presentation speed, suggesting that narrative processing reflects a dynamic interplay that is dependent on the structure of a narrative and processing constraints.

## ACKNOWLEDGEMENTS

This research was supported by the National Science Foundation award BCS-2335767 (LP) and a Research Catalyst Grant from New York University (LP).

We would like to thank David Poeppel for his helpful comments on the experiment design.

## REFERENCES

1. Davis MH. Chapter 44 - The Neurobiology of Lexical Access. In: Hickok G, Small SL, editors. Neurobiology of Language. San Diego: Academic Press; 2016. pp. 541–555. doi:10.1016/B978-0-12-407794-2.00044-4

2. Fedorenko E, Thompson-Schill SL. Reworking the language network. Trends Cogn Sci. 2014;18: 120–126. doi:10.1016/j.tics.2013.12.006

3. Friederici AD. Towards a neural basis of auditory sentence processing. Trends Cogn Sci. 2002;6: 78–84. doi:10.1016/S1364-6613(00)01839-8

4. Hagoort P, Indefrey P. The Neurobiology of Language Beyond Single Words. Annu Rev Neurosci. 2014;37: 347–362. doi:10.1146/annurev-neuro-071013-013847

5. Hickok G, Poeppel D. The cortical organization of speech processing. Nat Rev Neurosci. 2007;8: 393–402. doi:10.1038/nrn2113

6. Pylkkänen L. The neural basis of combinatory syntax and semantics. Science. 2019;366: 62–66. doi:10.1126/science.aax0050

7. Brennan JR, Pylkkänen L. MEG Evidence for Incremental Sentence Composition in the Anterior Temporal Lobe. Cogn Sci. 2017;41: 1515–1531. doi:10.1111/cogs.12445

8. Li J, Lai M, Pylkkänen L. Semantic composition in experimental and naturalistic paradigms. Imaging Neurosci. 2024;2: 1–17. doi:10.1162/imag_a_00072

9. Desbordes T, Lakretz Y, Chanoine V, Oquab M, Badier J-M, Trébuchon A, et al. Dimensionality and Ramping: Signatures of Sentence Integration in the Dynamics of Brains and Deep Language Models. J Neurosci. 2023;43: 5350–5364. doi:10.1523/JNEUROSCI.1163-22.2023

10. Fedorenko E, Scott TL, Brunner P, Coon WG, Pritchett B, Schalk G, et al. Neural correlate of the construction of sentence meaning. Proc Natl Acad Sci. 2016;113. doi:10.1073/pnas.1612132113

11. Pallier C, Devauchelle A-D, Dehaene S. Cortical representation of the constituent structure of sentences. Proc Natl Acad Sci. 2011;108: 2522–2527. doi:10.1073/pnas.1018711108

12. Bower GH, Morrow DG. Mental Models in Narrative Comprehension. Science. 1990;247: 44–48. doi:10.1126/science.2403694

13. Zwaan RA. From Words to Worlds: Twenty-Five Years of Advances in Situation Model Research. Curr Dir Psychol Sci. 2025;34: 287–292. doi:10.1177/09637214251326812

14. Zwaan RA, Radvansky GA. Situation models in language comprehension and memory. Psychol Bull. 1998;123: 162–185. doi:10.1037/0033-2909.123.2.162

15. Kurby CA, Zacks JM. Situation models in naturalistic comprehension. In: Willems RM, editor. Cognitive Neuroscience of Natural Language Use. Cambridge: Cambridge University Press; 2015. pp. 59–76. doi:10.1017/CBO9781107323667.004

16. Barnett A, Bellana B. Situation models and the default mode network. Curr Opin Behav Sci. 2025;66: 101593. doi:10.1016/j.cobeha.2025.101593

17. Xu J, Kemeny S, Park G, Frattali C, Braun A. Language in context: emergent features of word, sentence, and narrative comprehension. NeuroImage. 2005;25: 1002–1015. doi:10.1016/j.neuroimage.2004.12.013

18. Yarkoni T, Speer NK, Zacks JM. Neural substrates of narrative comprehension and memory. NeuroImage. 2008;41: 1408–1425. doi:10.1016/j.neuroimage.2008.03.062

19. Chang CHC, Nastase SA, Hasson U. Information flow across the cortical timescale hierarchy during narrative construction. Proc Natl Acad Sci. 2022;119: e2209307119. doi:10.1073/pnas.2209307119

20. Deniz F, Tseng C, Wehbe L, Tour TD la, Gallant JL. Semantic Representations during Language Comprehension Are Affected by Context. J Neurosci. 2023;43: 3144–3158. doi:10.1523/JNEUROSCI.2459-21.2023

21. Jain S, Huth A. Incorporating Context into Language Encoding Models for fMRI. Advances in Neural Information Processing Systems. Curran Associates, Inc.; 2018. Available: https://proceedings.neurips.cc/paper_files/paper/2018/hash/f471223d1a1614b58a7dc45c9d01df19-Abstract.html

22. Huth AG, De Heer WA, Griffiths TL, Theunissen FE, Gallant JL. Natural speech reveals the semantic maps that tile human cerebral cortex. Nature. 2016;532: 453–458. doi:10.1038/nature17637

23. Rubin G, Turano K. Reading without saccadic eye movements. Vision Res. 1992;32: 895–902. doi:10.1016/0042-6989(92)90032-E

24. Vagharchakian L, Dehaene-Lambertz G, Pallier C, Dehaene S. A Temporal Bottleneck in the Language Comprehension Network. J Neurosci. 2012;32: 9089–9102. doi:10.1523/JNEUROSCI.5685-11.2012

25. Wlotko EW, Federmeier KD. Time for prediction? The effect of presentation rate on predictive sentence comprehension during word-by-word reading. Cortex. 2015;68: 20–32. doi:10.1016/j.cortex.2015.03.014

26. Snell J, Grainger J. The sentence superiority effect revisited. Cognition. 2017;168: 217–221. doi:10.1016/j.cognition.2017.07.003

27. Fallon J, Pylkkänen L. Language at a glance: How our brains grasp linguistic structure from parallel visual input. Sci Adv. 2024;10: eadr9951. doi:10.1126/sciadv.adr9951

28. Flower N, Pylkkänen L. The Spatiotemporal Dynamics of Bottom–Up and Top–Down Processing during At-a-Glance Reading. J Neurosci. 2024;44. doi:10.1523/JNEUROSCI.0374-24.2024

29. Krogh S, Pylkkänen L. Manipulating syntax without taxing working memory: MEG correlates of syntactic dependencies in a verb-second language. Lang Cogn Neurosci. 2025;40: 1390–1413. doi:10.1080/23273798.2025.2549342

30. Li (李帛炫) B, Pylkkänen L. MEG investigation of adjective order preferences as a syntactic constraint. Cognition. 2026;273: 106536. doi:10.1016/j.cognition.2026.106536

31. Friederici AD. The Brain Basis of Language Processing: From Structure to Function. Physiol Rev. 2011;91: 1357–1392. doi:10.1152/physrev.00006.2011

32. Tarkiainen A, Helenius P, Hansen PC, Cornelissen PL, Salmelin R. Dynamics of letter string perception in the human occipitotemporal cortex. Brain. 1999;122: 2119–2132. doi:10.1093/brain/122.11.2119

33. Pylkkänen L. The neural basis of combinatory syntax and semantics. Science. 2019;366: 62–66. doi:10.1126/science.aax0050

34. Qi P, Zhang Y, Zhang Y, Bolton J, Manning CD. Stanza: A Python Natural Language Processing Toolkit for Many Human Languages. Proceedings of the 58th Annual Meeting of the Association for Computational Linguistics: System Demonstrations. Online: Association for Computational Linguistics; 2020. pp. 101–108. doi:10.18653/v1/2020.acl-demos.14

35. Culbertson G, Andersen E, Christiansen MH. Using Utterance Recall to Assess Second Language Proficiency. Lang Learn. 2020;70: 104–132. doi:10.1111/lang.12399

36. Peirce J, Gray JR, Simpson S, MacAskill M, Höchenberger R, Sogo H, et al. PsychoPy2: Experiments in behavior made easy. Behav Res Methods. 2019;51: 195–203. doi:10.3758/s13428-018-01193-y

37. Gramfort A, Luessi M, Larson E, Engemann DA, Strohmeier D, Brodbeck C, et al. MEG and EEG Data Analysis with MNE-Python. Front Neurosci. 2013;7: 1–13. doi:10.3389/fnins.2013.00267

38. Fischl B. FreeSurfer. NeuroImage. 2012;62: 774–781. doi:10.1016/j.neuroimage.2012.01.021

39. Maris E, Oostenveld R. Nonparametric statistical testing of EEG- and MEG-data. J Neurosci Methods. 2007;164: 177–190. doi:10.1016/j.jneumeth.2007.03.024

40. Youssofzadeh V, Conant L, Stout J, Ustine C, Humphries C, Gross WL, et al. Late dominance of the right hemisphere during narrative comprehension. NeuroImage. 2022;264: 119749. doi:10.1016/j.neuroimage.2022.119749

41. Gwilliams L, Lewis GA, Marantz A. Functional characterisation of letter-specific responses in time, space and current polarity using magnetoencephalography. NeuroImage. 2016;132: 320–333. doi:10.1016/j.neuroimage.2016.02.057

42. Brodbeck C, Das P, Brooks TL, Reddigari S, jpkulasingham. Eelbrain. Zenodo; 2023. doi:10.5281/zenodo.7951251

43. Huntenburg JM, Bazin P-L, Margulies DS. Large-Scale Gradients in Human Cortical Organization. Trends Cogn Sci. 2018;22: 21–31. doi:10.1016/j.tics.2017.11.002

44. Margulies DS, Ghosh SS, Goulas A, Falkiewicz M, Huntenburg JM, Langs G, et al. Situating the default-mode network along a principal gradient of macroscale cortical organization. Proc Natl Acad Sci. 2016;113: 12574–12579. doi:10.1073/pnas.1608282113

45. Hasson U, Yang E, Vallines I, Heeger DJ, Rubin N. A Hierarchy of Temporal Receptive Windows in Human Cortex. J Neurosci. 2008;28: 2539–2550. doi:10.1523/JNEUROSCI.5487-07.2008

46. Lerner Y, Honey CJ, Silbert LJ, Hasson U. Topographic Mapping of a Hierarchy of Temporal Receptive Windows Using a Narrated Story. J Neurosci. 2011;31: 2906–2915. doi:10.1523/JNEUROSCI.3684-10.2011

47. Aronson S, Adkins MS, Greene MR. Divergent roles of visual structure and conceptual meaning in scene detection and categorization. J Vis. 2025;25: 21. doi:10.1167/jov.25.14.21

48. Messi A-P, Pylkkanen L. Tracking Neural Correlates of Contextualized Meanings with Representational Similarity Analysis. J Neurosci. 2025;45. doi:10.1523/JNEUROSCI.0409-24.2025

49. Kim HJ, Lux BK, Lee E, Finn ES, Woo C-W. Brain decoding of spontaneous thought: Predictive modeling of self-relevance and valence using personal narratives. Proc Natl Acad Sci. 2024;121: e2401959121. doi:10.1073/pnas.2401959121

50. Dhankhar A, Wexler BE, Fulbright RK, Halwes T, Blamire AM, Shulman RG. Functional Magnetic Resonance Imaging Assessment of the Human Brain Auditory Cortex Response to Increasing Word Presentation Rates. J Neurophysiol. 1997;77: 476–483. doi:10.1152/jn.1997.77.1.476

51. Dupoux E, Green K. Perceptual adjustment to highly compressed speech: Effects of talker and rate changes. J Exp Psychol Hum Percept Perform. 1997;23: 914–927. doi:10.1037/0096-1523.23.3.914

52. Mehler J, Sebastian N, Altmann G, Dupoux E, Christophe A, Pallier C. Understanding Compressed Sentences: The Role of Rhythm and Meaning. Ann N Y Acad Sci. 1993;682: 272–282. doi:10.1111/j.1749-6632.1993.tb22975.x

53. Borges AFT, Giraud A-L, Mansvelder HD, Linkenkaer-Hansen K. Scale-Free Amplitude Modulation of Neuronal Oscillations Tracks Comprehension of Accelerated Speech. J Neurosci. 2018;38: 710–722. doi:10.1523/JNEUROSCI.1515-17.2017

54. Kutas M. Event-related brain potentials (ERPs) elicited during rapid serial visual presentation of congruous and incongruous sentences. Electroencephalogr Clin Neurophysiol Suppl. 1987;40: 406–411.

55. Kurby CA, Zacks JM. Situation models in naturalistic comprehension. In: Willems RM, editor. Cognitive Neuroscience of Natural Language Use. Cambridge: Cambridge University Press; 2015. pp. 59–76. doi:10.1017/CBO9781107323667.004

56. Long DL, Baynes K. Discourse Representation in the Two Cerebral Hemispheres. J Cogn Neurosci. 2002;14: 228–242. doi:10.1162/089892902317236867

57. Patel T, Morales M, Pickering MJ, Hoffman P. A common neural code for meaning in discourse production and comprehension. NeuroImage. 2023;279: 120295. doi:10.1016/j.neuroimage.2023.120295

58. Mason RA, Just MA. How the Brain Processes Causal Inferences in Text: A Theoretical Account of Generation and Integration Component Processes Utilizing Both Cerebral Hemispheres. Psychol Sci. 2004;15: 1–7. doi:10.1111/j.0963-7214.2004.01501001.x

59. Baumgarten TJ, Maniscalco B, Lee JL, Flounders MW, Abry P, He BJ. Neural integration underlying naturalistic prediction flexibly adapts to varying sensory input rate. Nat Commun. 2021;12: 2643. doi:10.1038/s41467-021-22632-z

60. Mandler G. Recognizing: The judgment of previous occurrence. Psychol Rev. 1980;87: 252–271. doi:10.1037/0033-295X.87.3.252

61. Yonelinas AP. Receiver-operating characteristics in recognition memory: Evidence for a dual-process model. J Exp Psychol Learn Mem Cogn. 1994;20: 1341–1354. doi:10.1037/0278-7393.20.6.1341

62. Boldini A, Russo R, Avons SE. One process is not enough! A speed-accuracy tradeoff study of recognition memory. Psychon Bull Rev. 2004;11: 353–361. doi:10.3758/BF03196582

63. Giglio L, Ostarek M, Weber K, Hagoort P. Commonalities and Asymmetries in the Neurobiological Infrastructure for Language Production and Comprehension. Cereb Cortex. 2022;32: 1405–1418. doi:10.1093/cercor/bhab287

64. Zacks JM, Speer NK, Swallow KM, Braver TS, Reynolds JR. Event perception: A mind-brain perspective. Psychol Bull. 2007;133: 273–293. doi:10.1037/0033-2909.133.2.273

65. Speer NK, Zacks JM, Reynolds JR. Human brain activity time-locked to narrative event boundaries. Psychol Sci. 2007;18: 449–455. doi:10.1111/j.1467-9280.2007.01920.x

66. Chen J, Bornstein AM. The causal structure and computational value of narratives. Trends Cogn Sci. 2024;28: 769–781. doi:10.1016/j.tics.2024.04.003

67. Song H, Park B, Park H, Shim WM. Cognitive and Neural State Dynamics of Narrative Comprehension. J Neurosci. 2021;41: 8972–8990. doi:10.1523/JNEUROSCI.0037-21.2021

68. Just MA, Carpenter PA. A capacity theory of comprehension: Individual differences in working memory. Psychol Rev. 1992;99: 122–149. doi:10.1037/0033-295X.99.1.122

